# Engineering IspH for Enhanced Terpenoid Yield: Computational and Molecular Dynamics Studies

**DOI:** 10.1101/2025.04.07.643449

**Authors:** Ashish Runthala, Sivudu Macherla, Venkatramanan Varadharajan, Silambarasan Tamil Selvan, Manmohan Sharma, Nameet Kaur, Suresh Chandra Phulara, Noor Ahmad Shaik, Hazem K Ghneim

## Abstract

Terpenoids play a vital role in pharmaceuticals, biofuels, and various industries, and they are produced through the methylerythritol phosphate (MEP) pathway, with IspH catalyzing the final step. This study explores IspH homologs in Bacillus and other bacteria to aid in enzyme engineering. Sequence analysis showed a conservation range of 46-79%. Phylogenetic analysis (log-likelihood -71,581.08, bootstrap 80-100) validated the evolutionary relationships. Structural modeling revealed conserved functional motifs. Molecular docking indicated HMBPP binding affinities between -4.9 and -6.4 kcal/mol, while molecular dynamics simulations confirmed the stability of the complex (RMSD 2.02-3.45 Å, Rg 21.56-22.06 Å, with a decrease in SASA upon binding). Significant hydrogen bonds were noted with His131, Asn227, and Ser271. This thorough analysis of IspH conservation, structure, and stability, along with co-evolving residue networks, underscores the potential of Bacillus IspH variants for improved terpenoid production and lays the groundwork for targeted enzyme engineering.

## 1. Introduction

With more than 70,000 unique compounds indexed in the molecular dictionary of natural products, terpenes and their various derivatives are one of the largest and structurally most diverse categories of natural chemicals^1^. These molecules serve a variety of functions, ranging from protein prenylation, mediating antagonistic/symbiotic interactions between organisms to electron transfer or contribution to membrane fluidity^2,3^. Due to their vast functional and structural variability, terpenoids have been used for a variety of applications, including pharmaceuticals, biofuels, tastes, and scents^4–7^. Although there are still many of these compounds that are taken from the natural sources, including plants, this approach frequently suffers from seasonal and geographic changes in supply and quality. One excellent illustration of poor yields or perhaps insufficient plant material is the potent anticancer drug Taxol (paclitaxel), which was found in the bark of adult pacific yew trees^8^.

Biotechnological production of chemicals, especially terpenoids, has gained significant interest over the last two decades, driven by a shift toward sustainable manufacturing^9–12^. The study has been focused on using the most genetically traceable and tractable microbial host E. coli^13–14^, because certain aspects, including genetic engineering, characterisation, reliability, and independence of biological modules are still fraught with uncertainty. To overcome such limitations, several other microbial hosts, both from prokaryotic and eukaryotic communities, have been explored in recent years. The eukaryotic host, *Saccharomyces cerevisiae*, has contributed substantially towards the production of higher-order terpenoids^15^; however, *Bacillus subtilis* has been actively favoured for biosynthesizing the lower-order terpenoids^16,17^. While isopentenyl diphosphate (IPP) and dimethylallyl diphosphate (DMAPP) are endogenously synthesized via the mevalonate pathway in Eukarya, Archaea, and some bacteria^18–21^, recent research has identified an alternative methylerythritol 4-phosphate (MEP) pathway for their biosynthesis in plant plastids and most eubacteria^21–23^. Condensation of two or more of these molecules leads to the formation of the larger prenyl diphosphate compounds like geranyl diphosphate (GPP), farnesyl diphosphate (FPP), and geranylgeranyl diphosphate (GGPP). To date, the intermediates and enzymes of the MEP pathway have all been identified ^24^, and this extensive knowledgebase has been utilized to produce several valuable hemiterpenoids (C5), monoterpenoids (C10), sesquiterpenoids (C15), diterpenoids (C20), sesterterpenoids (C25), triterpenoids (C30), and tetraterpenoid (C40) molecules^25–32^.

The biosynthesis of isopentenols (prenol and isoprenol), hemiterpenoid-based alcohols with potential applications as nutraceuticals has recently been explored using the *B. subtilis* MEP pathway^33^. While the catalytic activity of pyrophosphatases on DMAPP synthesizes prenol, hydrolysis of IPP by pyrophosphatases leads to the synthesis of isoprenol, as shown in Figure1. However, wild-type *Bacillus subtilis* produces only a small amount of isoprenol, likely due to the inherently low level of precursor pyrophosphates and the low catalytic activity of IspH. Hence, the production of isopentenols heavily depends on IPP and DMAPP flux, which can be optimized by modulating the activity of the IspH enzyme. In an effort to enhance productivity, the MEP pathway has even been integrated with the Glycolysis pathway in *Escherichia coli* to increase yields of downstream molecules^34^.

**Figure 1:**
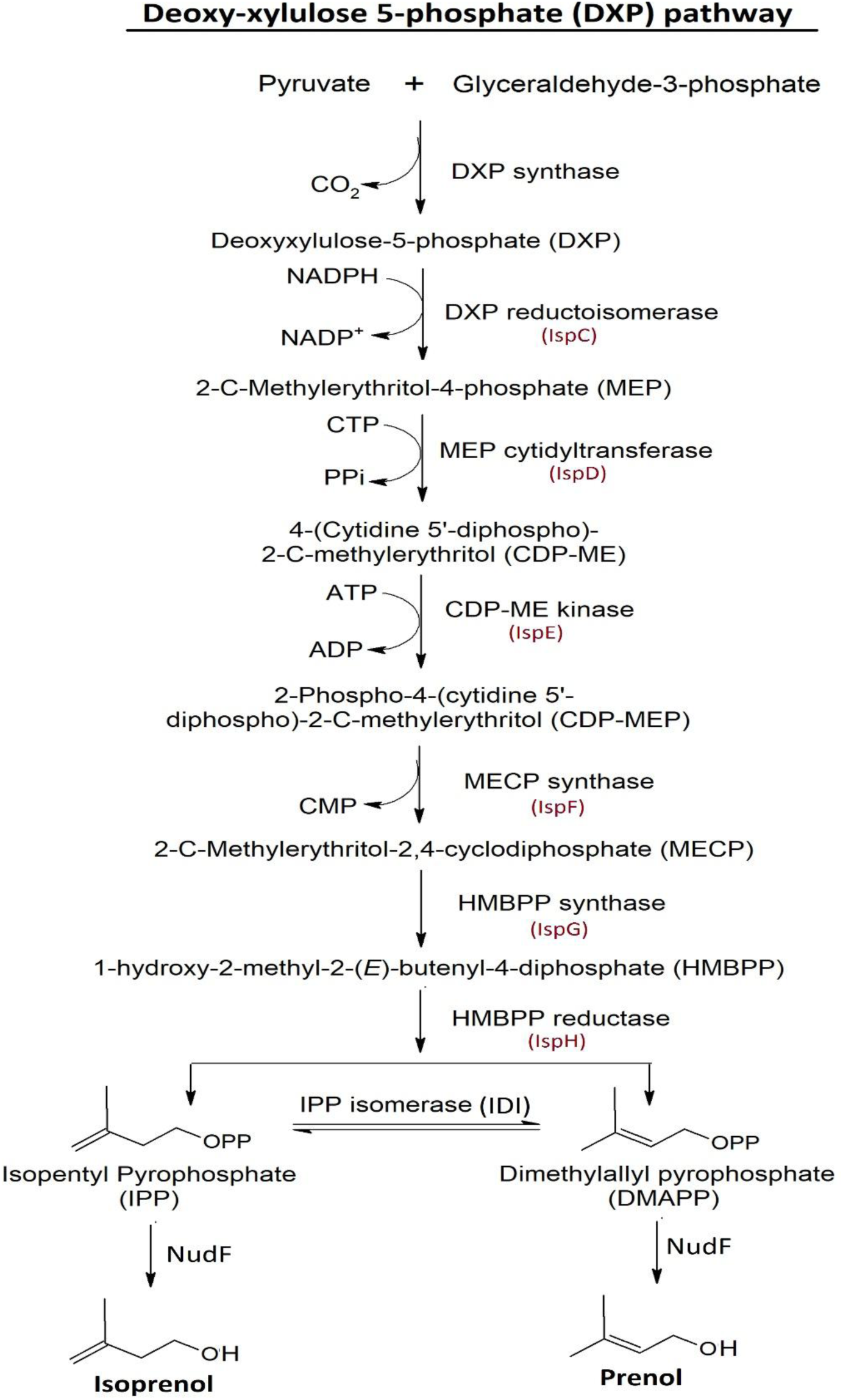
MEP pathway: This metabolic process produces IPP and DMAPP, which serve as the 5-carbon building blocks for isoprenoid biosynthesis. Glyceraldehyde 3-phophate and pyruvate are condensed using DXS, and IPP and DMAPP are produced in six more steps.

IspH, the terminal enzyme of the MEP pathway, converts 1-hydroxy-2-methyl-2-(E)-butenyl-4-diphosphate (HMBPP) into dimethylallyl pyrophosphate (DMAPP) and isopentenyl pyrophosphate (IPP) in a 1:5 ratio, directing carbon flux toward terpenoid synthesis^35^. Its expression level significantly influences pathway efficiency, as excess HMBPP can lead to cellular toxicity and reduced terpenoid yield^36^. IspH expression is thus tightly regulated to control HMBPP toxicity without surpassing optimal catalytic activity, contrasting with enzymes like DXS and DXR, which are overexpressed to boost yields^37^. Synthetic operons have even enabled stable multigene expression for terpenoid synthesis in *Bacillus subtilis*^38^, yet further understanding of IspH’s catalytic mechanisms is needed for optimized IPP and DMAPP production, especially in prokaryotic systems^39,40^. Structural studies reveal that IspH houses a central [4Fe-4S] cluster essential for catalysis, though an Fe3S4 variant has also been reported^41,42^. This cluster facilitates electron transfer and protonation events necessary for HMBPP reduction, with a structural shift from an open to closed conformation upon substrate binding that enhances catalytic efficiency^43,44^. Molecular simulations show domain flexibility, creating adaptable substrate binding pockets.^43^ Current inhibitors exhibit limited binding efficacy to IspH’s iron site, signalling a need for the detailed understanding of IspH structure for several microorganism^44–45^.

In light of the need for comprehensive analysis on the evolutionary coupling, interaction network, and phylogenetic promiscuity of IspH, our study aims to thoroughly investigate the non-redundant sequence datasets from *Bacillus subtilis*, *Brevibacillus brevis*, and other bacterial species. We hypothesize that the study shall provide us logical framework and help us investigate the physiological function of IspH for the heterologous production of more complex terpenoids in *B. subtilis*.

## 2. Materials and Methods

### 2.1 Retrieving the functionally correct IspH sequences

Screening UniProt^46^ and Expasy (EC: 1.17.7.4), a gigantic sequence dataset of 25568 entries is constructed. As per the length range of 297-335 for structurally resolved and well-annotated protein IspH structures, a relaxed filter of 275-350 is used to select the functionally similar sequences, and the outlier sequence lengths are discarded. It allowed us to construct a dataset of 399 sequences, of which only 34 and 12 belong to *Brevibacillus brevis*47 and *B. subtilis*168 strains, respectively. To further analyze the evolutionary conservation of these datasets in contrast to the other species, the remaining entries are thus considered as set3, comprising 353 sequences belonging to other species.

### 2.2 Modelling the representative sequences of the three sets and affirming their functional nature

As nature reduces the protein sequence length in evolution to pack the information with a smaller number of amino acids, the smallest sequence from each dataset is modelled to decipher its structural details.^47^ As demonstrated earlier^48–49^, the smallest proteins C0ZL73, P54473, and A5IME1, more representative of the functionally crucial residues/motifs for the datasets set1-set3, are selected as the representative proteins, orderly belong to the species *B. brevis*47, *B. subtilis*168, and *Thermotoga petrophila,* for the downstream analysis. As similarities within the relatively shorter regions of a longer sequence causes alignment error^49^, consideration of smaller sequences should make the further processing more accurate. To precisely map the evolutionarily retained substructures, the tertiary structures are predicted using Alphafold^50^. For validating the functional similarity of these proteins, their top-3 motifs are further identified by the MEME program (Version 5.4.1, https://meme-suite.org/meme/tools/meme (accessed on February 28, 2024) ^51^.

### 2.3. Tracking the evolutionary variation

To analyze the functional divergence across various sequence datasets for all the biochemically crucial positions, a two step-methodology is utilized. Sequence datasets are aligned using the HMM-based ClustalO module of HHPred^52^. By screening the query HMM against the pre-compiled sequence profile database of HHPred, this sensitive homology screening method constructs biologically optimal alignments, enhancing the accuracy of downstream analysis^48, 53, 54^. Sequence conservation is further analyzed using the Consurf to map the positional variations across the aligned residues in correlation with their topologically-buried/-exposed nature^55^. The alignments are fed into the IQ-Tree server, and the methodology is implemented using 1000 bootstrapped alignments with a minimum correlation coefficient of 0.99 for 1000 iterations to enhance prediction accuracy^56^. The constructed consensus tree is visualized using ITOL to accurately map the sequence divergence across the sets^57^.

### 2.4. Tracing the evolutionarily coupled network

Maintaining the functionally crucial interaction network against the different pathway proteins requires a strongly conserved molecular coupling of residues. Hence, the coevolved residue network is traced for all the constructed sequence alignments of the three datasets. For attaining a higher accuracy of prediction, the recently benchmarked Leri server is utilized^58^, and the results are compared using the computed scores.

### 2.5. Building the IspH-HMBPP complexes

To screen the interaction of the modelled proteins against HMBPP, their key binding site is computationally localized using CASTp server^59^. For an ease of downstream analysis, P54473 and A5IMEI models are transposed over the reference molecule C0ZL73. Moreover, for removing the water molecules and other inhibitors/heteroatoms, the full-length modelled protein structures are prepared using the Discovery studio visualizer, as usually done^60^. Docking simulations were performed using PyRx 0.8 (Autodock Vina, Scripps Research Institute, La Jolla, CA) to predict the binding mode of HMBPP within C0ZL73. PyRx facilitated the conversion of protein and ligand structures to the PDBQT format and employed Openbabel for energy minimization. For docking HMBPP in all the defined site of all the three proteins, TYR13-OH (-2.872, 4.565, -12.311), located in the central cavity of C0ZL73, is designated as the gridcenter to define the gridbox of 25Åx25Åx25Å.

### 2.6. Molecular dynamics simulation

To explore the thermodynamic characteristics and stability of the anticipated protein-ligand complexes, molecular dynamics (MD) simulations are performed using the Desmond v7.2 module of the Maestro v12.5 suite^61^. Removing water molecules beyond 5Å, assigning bond ordering, adding hydrogens, and disulfide bonds, the structures are arranged in an orthorhombic box with 10Å periodic boundaries and solvated using the simple point charge (SPC) water model^62^. The molecular systems are neutralized using 0.15M NaCl and energetically relaxed using the OPLS3e force field^63^, with the SHAKE algorithm constraining water molecule geometries and the bond lengths and angles of heavy atoms^64^. Periodic boundary conditions and the particle mesh Ewald method manage long-range electrostatic interactions^65^. Simulations are conducted in the NPT ensemble, maintaining a temperature of 300K and pressure of 1.01325bar for 100 ns, with data collection intervals of 100 ps. Temperature and pressure are controlled via the Nose-Hoover thermostat^66^, and Martyna-Tobias-Klein^67^ methods, respectively. The RESPA integrator with a 2.0fs timestep is used for bonded and non-bonded interactions^68^. For screening the IspH interaction against HMBPP for the three representative sequences, the simulations are lastly analyzed using four key matrices: root mean square deviation (RMSD), root mean square fluctuation (RMSF), radius of gyration (Rg) and solvent accessible surface area (SASA). The intermolecular interactions between the ligand and protein, particularly the formation of hydrogen bonds throughout the simulation, are analyzed using the simulation interaction diagram package.

## 3. Results and Discussion

### 3.1 Building the IspH sequence datasets

Screening UniProt and the enzyme category database of Expasy (EC: 1.17.7.4), a gigantic sequence dataset of 25568 entries is constructed. As per the length range of 297-335 for structurally resolved and well-annotated protein IspH structures, a relaxed filter of 275-350 is used to select the functionally similar sequences and discard the outlier sequences. This enabled us to compile a dataset of 399 sequences, with only 34 and 12 specifically belonging to the *B. brevis* 47 and *B. subtilis* 168 strains, respectively. The derived datasets will allow us to extract the crucial cellular functional details, aiding us to develop a deeper understanding about this important cellular biofactory^69^. To further assess the evolutionary conservation of these datasets in comparison to other species, the remaining entries are designated as Set3, consisting of 353 sequences from various species.

### 3.2 Modelling the representative protein sequences

To structurally analyze the representative protein sequences C0ZL73, P54473, and A5IME1 of the three datasets set1-3, their conformation is predicted through Alphafold^50^. The structural superimposition of these models indicates a strong topological similarity across the chains (Figure 2A), as expected. The analysis further reveals that P54473 has an RMSD of 0.711 versus C0ZL73 for 311 overlapping atom pairs, C0ZL73 has an RMSD of 4.149 when compared to A5IME1, and P54473 shows an RMSD of 4.259 for 272 atom pairs against C0ZL73 (Figure 2B). Observing the resultant figure, the topological displacement of loops and a few secondary structures can be observed in these models, indicating that their biologically active core must be highly conserved, as shown by the low RMSD of 0.545, 1.17, and 1.186 for the 299 (96.141%), 126 (40.127%), and 134 (48.727%) tightly overlapped pairs within a distance deviation of 5Å.

**Figure 2:**
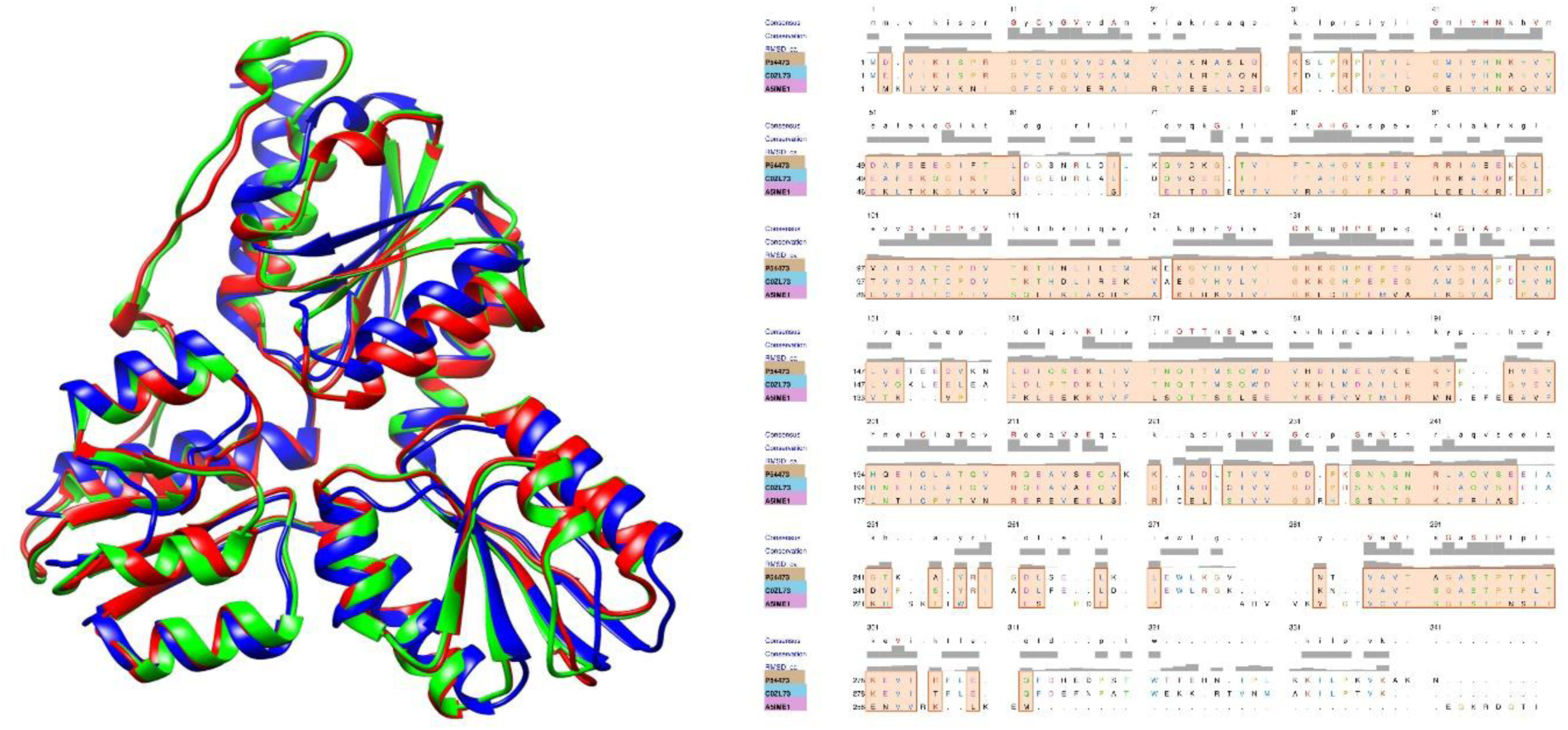
Chimera-based (A) structural superimposition of representative proteins C0ZL73 (green), P54473 (red), and AM5IME1 (blue), and their (B) structural alignment, indicating a high Cα-RMSD score is found to be very high for some residues.

As expected, the two *Bacillus* representatives for set1 and set2 show a colocalized set of three motifs, although the set3 representative encodes only a single motif (Figure 3). It affirms the credibility of the sequence datasets and the results. The Bacillus representatives for set1 and set2 have three colocalized motifs, while set3 has one, confirming the dataset’s credibility and aligning with the recently observed downstream flux variation^69^.

**Figure 3:**
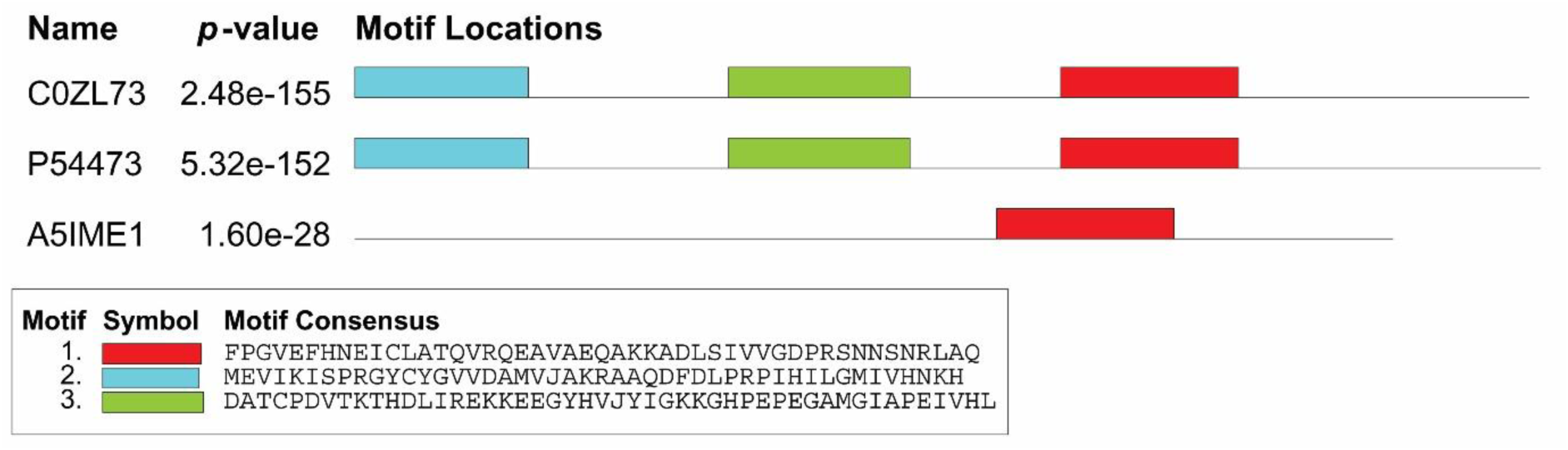
Conserved motif analysis of the three representatives.

### 3.3. Tracking the evolutionary variation

As demonstrated earlier,^48–53^ the HMM-profile of the sequence datasets is constructed using HHPred and is fed to Consurf.^55^ Contrary to set3 sequences, which only encode 46.178% of conserved residues, with a conservation score of at least 7, the set1 and set2 datasets show a relatively higher percentage of 50% and 78.98% conserved residues (Figure 4A-C). However, using the color gradient strategy (Figure 4D), it is observed that set1-set3 datasets orderly show 15.28%, 7.32% and 14.96% highly variable residues with a conservation score lesser than 4. Moreover, plotting these conservation scores, it is observed that set1 and set3 share an almost similar trend of residue conservation (Figure 4E), unlike set2, indicating that the IspH enzyme is remarkably conserved in *B. subtilis168*. Further, in line with the earlier research^42, 70–72^, and using several mutants for these positions, their roles behind the promiscuous catalytic activities of their several IspH mutants have been recently investigated^73^. The strictly conserved residue mutations H131N and E133Q of *B. subtilis sp. N16-5*, as well as H124N and E126N of *E.coli*, have been shown to result in the loss of isoamylene synthesis activity, although these IspH variations could still synthesize isoprene^74^. Moreover, mutating the three completely conserved cysteines have also been shown to decrease the *E.coli* IspH activity 70000 times^74^, proving the biological significance of conserved residues in IspH [42, 70, 75], as rightly shown by Figure 4E^42,70,75^.

**Figure 4:**
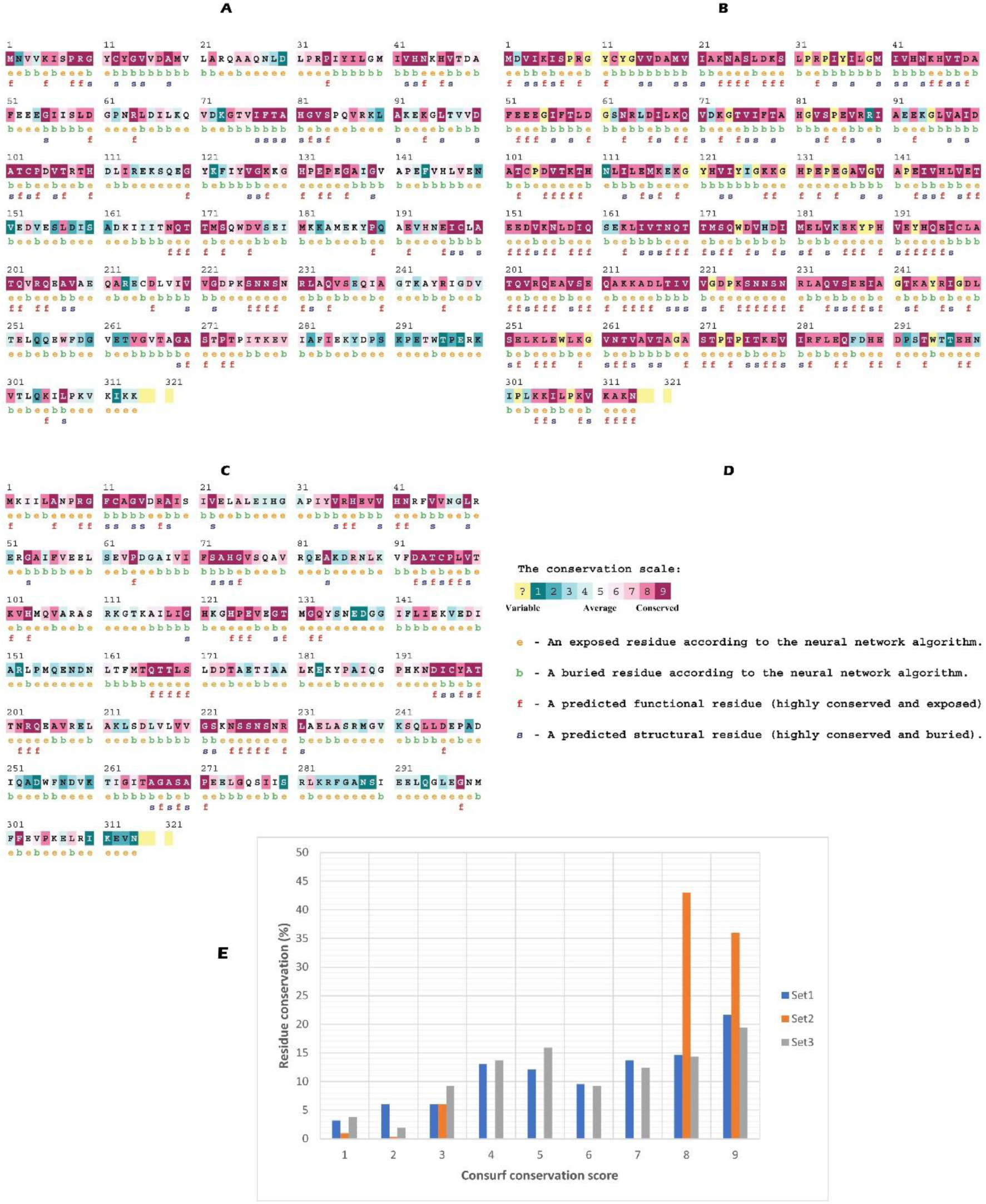
Consurf result for the three datasets (A) set1, (B) set2, and (C) set3. The conservation scale of the residues is illustrated using a (D) color gradient, besides indicating their structurally exposed/buried nature. (E) Sequence conservation plot of the three datasets.

Besides the assessment of sequence conservation for the three datasets, their overall evolutionary relationship is analysed using IQ-tree^56^. The analysis involved a total of 399 proteins, derived using the statistical correlation coefficient of 0.99, as shown earlier^76^. For the considered alignment, the number of invariant sites is found to be 10.8607%. The resultant consensus phylogenetic tree shows the log-likelihood score of -71581.0847 (Figure 5), and its credibility is also justified by a substantially higher bootstrap score of all the branches, ranging from 80-100^76^. Visualizing it using ITOL^57^, a total of 4 clades are observed for the three sequence datasets set1-set3, orderly represented as blue, red and green. While clades 3 and 4 have set1 and set3 proteins, clades 2 and 1 only encompass the set2 and set3 entries respectively. In comparison to the average mutual sequence similarity of 63.531±24.636 of the set1 entries, set2 and set3 orderly shows the scores of 97.827±1.025, and 46.429±16.501. It explains the tight clustering of set2 clade compared to set1 and set3, where a lower mutual similarity score leads to an increased number of their cladistic bifurcations.

**Figure 5:**
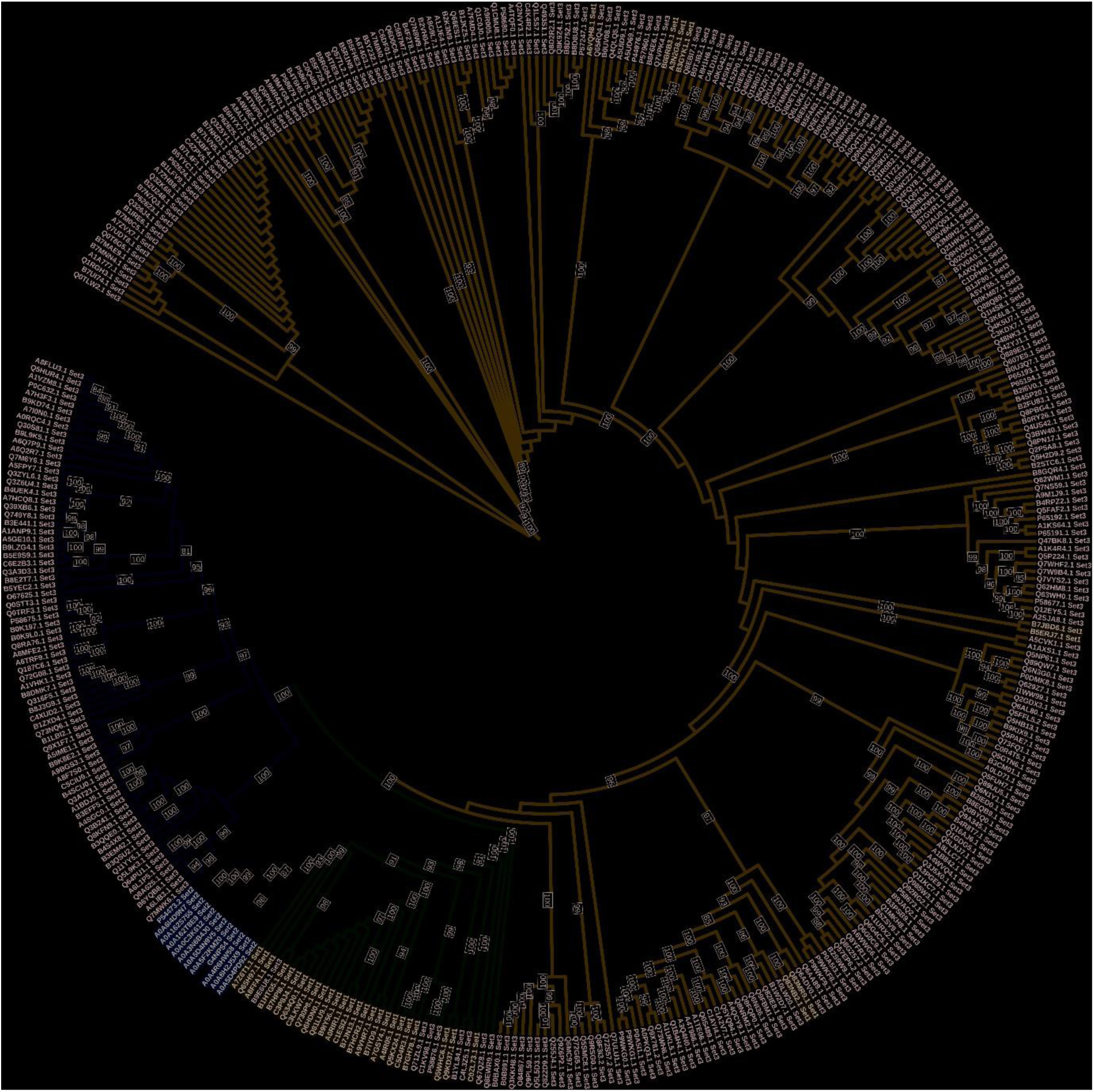
Phylogenetic tree of the three datasets, showing four different clades for the three datasets set1-set3, represented as blue, red, and green respectively.

### 3.4. Tracing the evolutionarily coupled network

Evolutionarily coupled residue pairs define the special set of residues that coevolve within a protein and govern its overall structure/function. It implies that mutations of the first residue are topologically compensated by the concomitant mutations at the second position^77^. Such functionally important residue-couples might be easily traced by examining the pattern(s) of sequence conservation and co-variation for homologous proteins of several species, and this could be helpful in functionally improving the stability, activity, or selectivity of a protein sequence^77,78^.

Building upon the established reliability of evolutionary coupling for protein folding^79^, our analysis of sets 1-3 reveals functionally crucial evolutionary signatures within distinct residue communities (Figure 6). Unlike sets 2 and 3, set1 shows two clusters: a 7-residue group (K74, L80, I83, K126, I132, E157, A174) and a 14-residue group (M1, K2, E86, E121, T152, N211, S251, L264, E266, M267, E268, G269, K270, R271), with E86 bridging both. These observations suggest two key points. First, the *Bacillus* strain exhibits a higher degree of evolutionary coupling compared to *B. subtilis* (Figure 6A), potentially reflecting the conserved linkage between key residues noted in recent research on horizontal gene transfer in *B. subtilis*^80^. Second, while most residues show some variation across the protein family (Figure 6B), the identified evolutionary couplings likely hold functional importance due to their non-absolute conservation. Identifying the functionally crucial residues, this experimental step offers a valuable resource to guide the development of targeted mutagenesis experiments for bacterial IspH homologues.

**Figure 6:**
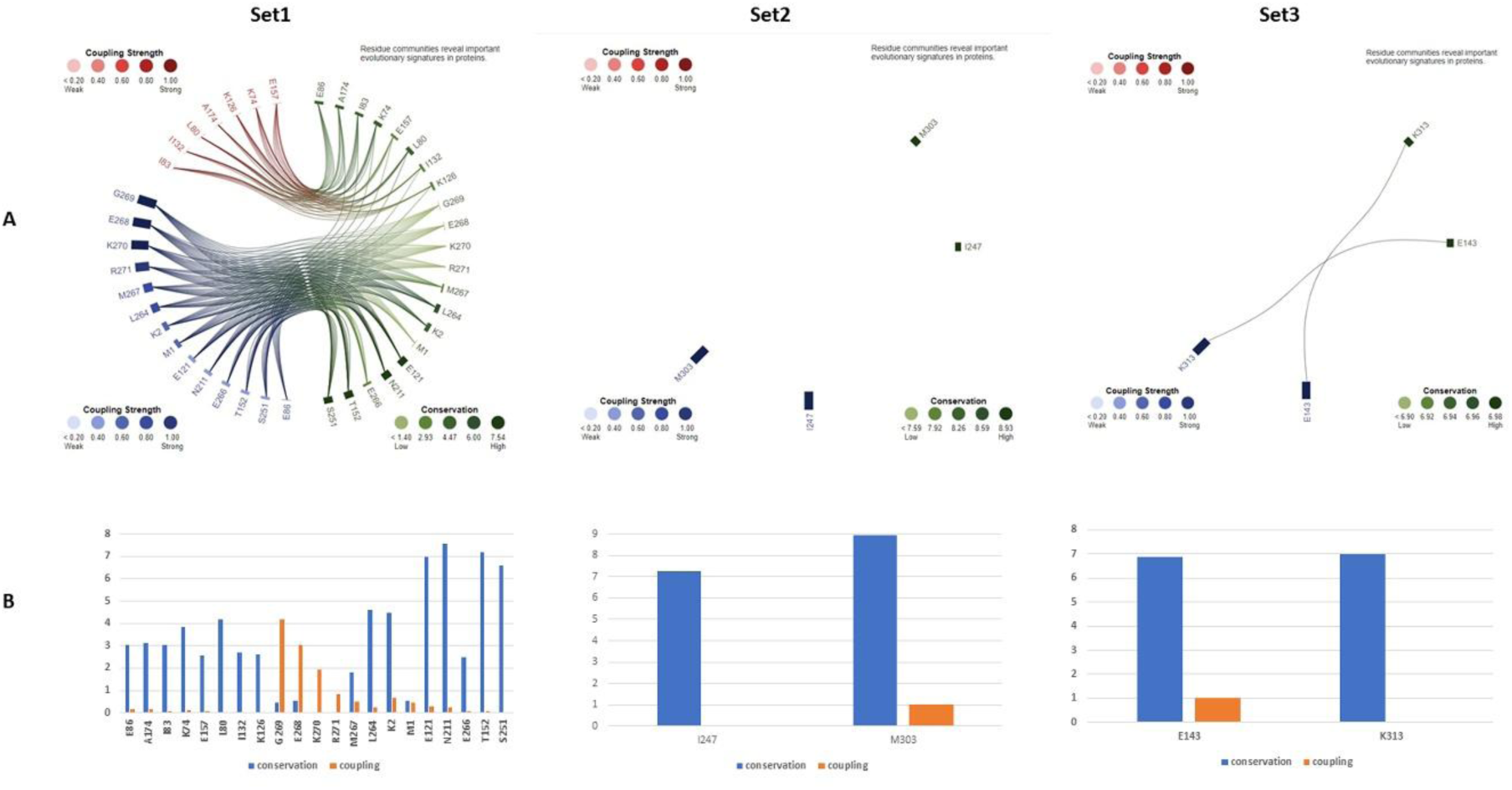
(A) Evolutionarily coupled residue pairs for the three datasets, and (B) conservation score for these residues.

### 3.5. Building the IspH-HMBPP complexes

Using the default probe radius of 1.4Å within CASTp, the most voluminous active site is screened for the modelled representative IspH proteins. Superimposition of these three proteins using Chimera highlights a very high topological similarity, indicating a substantial colocation of their active sites (Figure 7). For C0ZL73, the cavity, encompassed by 23 residues (CYS12, TYR13, GLY14, VAL15, VAL42, HIS43, ALA80, HIS81, CYS103, ASP105, VAL106, HIS131, GLU133, THR170, THR171, CYS198, ALA200, THR201, SER226, ASN227, ASN228, ALA270, and SER271) shows a surface area and volume of 172.79Å^2^, and 85.171Å^3^ respectively. The protein C0ZL73 and HMBPP exhibit a favorable interaction energy of -6.4 kcal/mol when docked together, suggesting that HMBPP may bind firmly within the active site. Likewise, against the same ligand, P54473 and A5IME1 orderly exhibit the interaction energies of -6.5, and -4.9Kcal/mol, indicating a substantial variation in the active site topology of these proteins.

**Figure 7:**
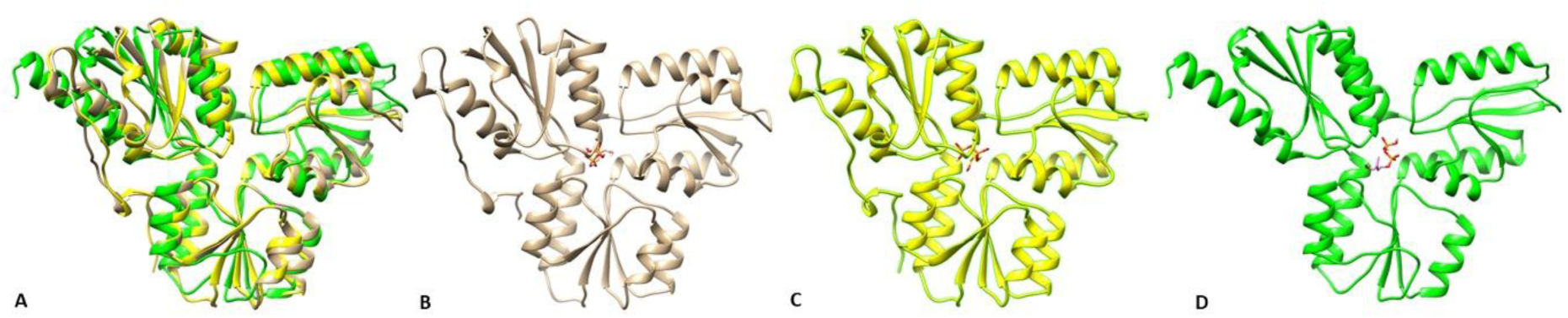
(A) Structural superimposition, and the docked complexes of the proteins (B) C0ZL73 (colorless), (C) P54473 (yellow), (D) A5IME1 (green) with the ligand HMBPP, highlighting a robust topological conservation with a pairwise Cα-RMSD ranging from 0.545 to 2.105.

### 3.6. MD simulation

To assess the stability of the predicted protein structures in their apo forms and HMBPP-bound complexes, molecular simulations were carried out for 100 ns. The trajectories are analyzed over the simulation time using several scoring measures viz. RMSD, RMSF, Rg, and SASA. The residues involved in various interactions, including hydrogen bonds, hydrophobic interactions, water bridges, and salt bridges, are analyzed along with the fractions of time these interactions persisted during the simulation

#### 3.6.1 RMSD, RMSF, Rg, and SASA analysis

The trajectories of the protein and ligand relative to the protein are analyzed over the simulation time, with comparisons made to the reference frame at 0 ns to compute RMSD, and RMSF scores. RMSD quantifies the average deviation of atoms in a simulated structure compared to a reference, providing insights into overall structural changes throughout simulation^81^. Conversely, RMSF measures the average deviation of each atom from its average position over the trajectory, offering information about local flexibility and mobility within the molecule^82^. In the case of protein C0ZL73 (Figure 8A), the RMSD value increases immediately at the start of the simulation, reaching a peak of 2.80 Å at 27.2ns. As the simulation progresses, the RMSD value decreases until 58ns, then gradually increases, peaking at 3.31 Å at 99ns. The average RMSD value observed after equilibrium, attained at 27ns, is 2.40 Å. When C0ZL73 is complexed with HMBPP, an initial rise in RMSD is observed until 3 ns, followed by a gradual decrease to 1.28Å at 26ns. Subsequently, the RMSD value increases, reaching a maximum of 3.35 Å at 49.6ns. Afterward, the RMSD value fluctuates and maintains equilibrium until 100ns. The average RMSD value observed for the complex is 2.02 Å, which is lower than the average RMSD observed for the apo-form of C0ZL73. For P54473 in its apo form (Figure 8B), the RMSD value increases gradually from the start of the simulation, reaching a peak 2.89Å at 10.3ns, and subsequently, the RMSD curve stabilizes, with an average RMSD value of 2.60Å. For its HMBPP-complex, an immediate spike in RMSD is observed at the beginning of the simulation, reaching a maximum of 2.75 Å at 0.9ns. The RMSD score then decreases steadily until 68ns, reaching an equilibrium with an average RMSD value of 1.76 Å. Morover, A5IME1 exhibits a peak RMSD value of 4Å at 18.2ns (Figure 8C), after which the RMSD curve stabilizes with an average RMSD of 3.45Å. The HMBPP complex exhibits a similar RMSD fluctuation pattern, with an average RMSD of 4.82 Å after equilibrium and a maximum RMSD of 5.5 Å at 23.3 ns. Typically, a high RMSD value indicates significant conformational change, and based on the observed average RMSD values, it can be deduced that proteins C0ZL73 and P54473, as well as their HMBPP-complexes, are more stable (RMSD < 3 Å) than A5IME1-complex, as also shown earlier^83^. The RMSF map in Figure 8D illustrates the conformational dynamics experienced by each residue in C0ZL73 and its complex with HMBPP. Notable peak RMS fluctuations in C0ZL73 are observed in residues such as Gly61, Glu62, Glu155, Ala156, Leu157, Pro160, Ala304, Val310, and Lys311. Upon binding to HMBPP, the C0ZL73-HMBPP complex exhibits a significant increase in RMS fluctuations in several residues, including Pro160, Phe290, Ala293, Thr294, and Lys311. Intriguingly, none of these residues directly interact with the ligand, suggesting that the observed fluctuations might be attributed to allosteric effects or long-range conformational changes induced by ligand binding. These findings highlight the complex interplay between ligand binding and protein flexibility, emphasizing the potential role of distal residues in modulating the overall conformational landscape of the protein. Significant RMS changes are detected in residues Gln160, His289, Glu290, Asp291, Pro292, Ser293, Asn300, Ile301, Pro302, and Leu303 of P54473 (Figure 8E). A comparable RMSF profile is observed for its HMBPP-complex, with significant fluctuations in RMSF values in areas such as Glu118-Gly120 and His178-Pro189. During the simulation, no residues with significant RMS fluctuations make contact with the ligand, as was the case with the C0ZL73 complex. The RMSF plots for A5IME1 and its complex with HMBPP are shown in Figures 8C and 8F, respectively. Increased RMSF values are observed for the residues Ile59, Thr60, Asp61, Lys107, Glu108, Asp272, Gly273, Thr274, and Ile275. A5IME1-HMBPP complex also show a similar RMSF pattern, with shifts in the region from Lys7 to Asn41, and 8 residues viz. Asn8, Cys12, Phe13, Gly14, Val15, Ile83, Val39, and His40 are observed to interact with HMBPP. Most regions that exhibit RMSF spikes are those connecting secondary structures such as helices and sheets, which lack stabilizing binding interactions^84,85^. Thus, for proteins C0ZL73 and P54473, the ligand does not engage with the flexible residues, resulting in extremely stable protein-ligand interactions. In contrast, the ligand’s interaction to A5IME1 causes enhanced flexibility in particular residues, as evidenced by elevated RMSF values. This greater flexibility could lead to less stable interactions between A5IME1 and HMBPP.

**Figure 8:**
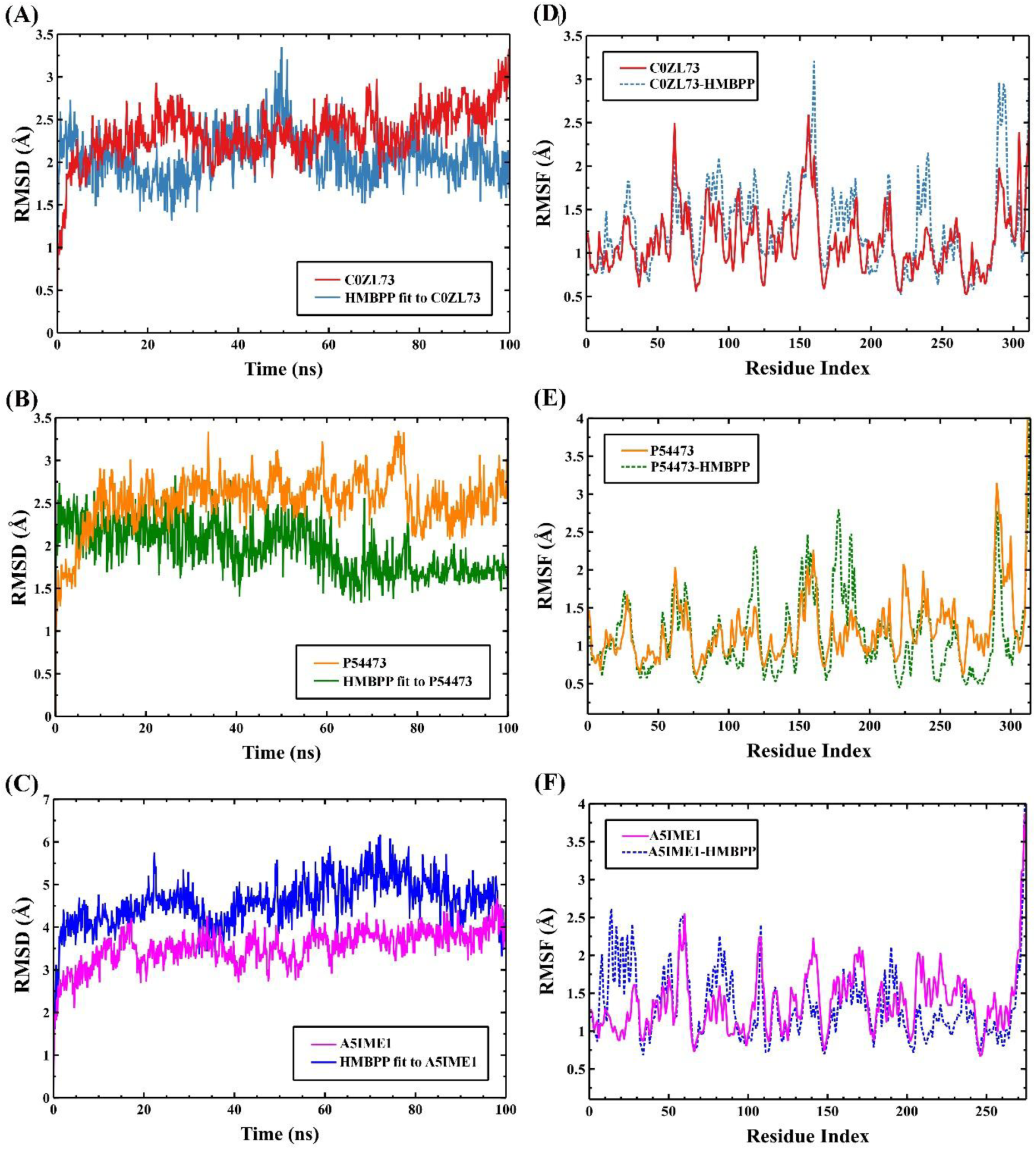
The RMSD and RMSF plots for the 100 ns MD simulation of the protein and protein-ligand complexes. C0ZL73-HMBPP (A, D), P54473-HMBPP (B, E), and A5IME1-HMBPP (C, F).

In addition to RMSD and RMSF, the radius of gyration (Rg) and solvent-accessible surface area (SASA) are critical metrics for assessing the simulation trajectory and gaining precise insights into the structural characteristics and stability of biological molecules. Rg measures the compactness of a molecular structure, providing insights into the protein’s folding state and overall stability, with consistent Rg values indicating structural stability^86^. SASA estimates the surface area of a biomolecule that is accessible to solvents^87^. The Rg and SASA plots for the proteins and their complexes with HMBPP are depicted in Figure 9. Initially, the Rg values exhibit a gradual increase (Figures 9A-C). Subsequently, the Rg values for C0ZL73 and A5IME1 stabilize, while P54473 shows a continued increase after 69 ns. The average Rg values for C0ZL73, P54473, and A5IME1 are 22.11Å, 21.77Å, and 22.13Å, respectively. However, upon forming complexes with HMBPP, a slight reduction in the average Rg values is observed, with the corresponding values being 22.06Å, 21.56Å, and 21.99Å, respectively.

**Figure 9:**
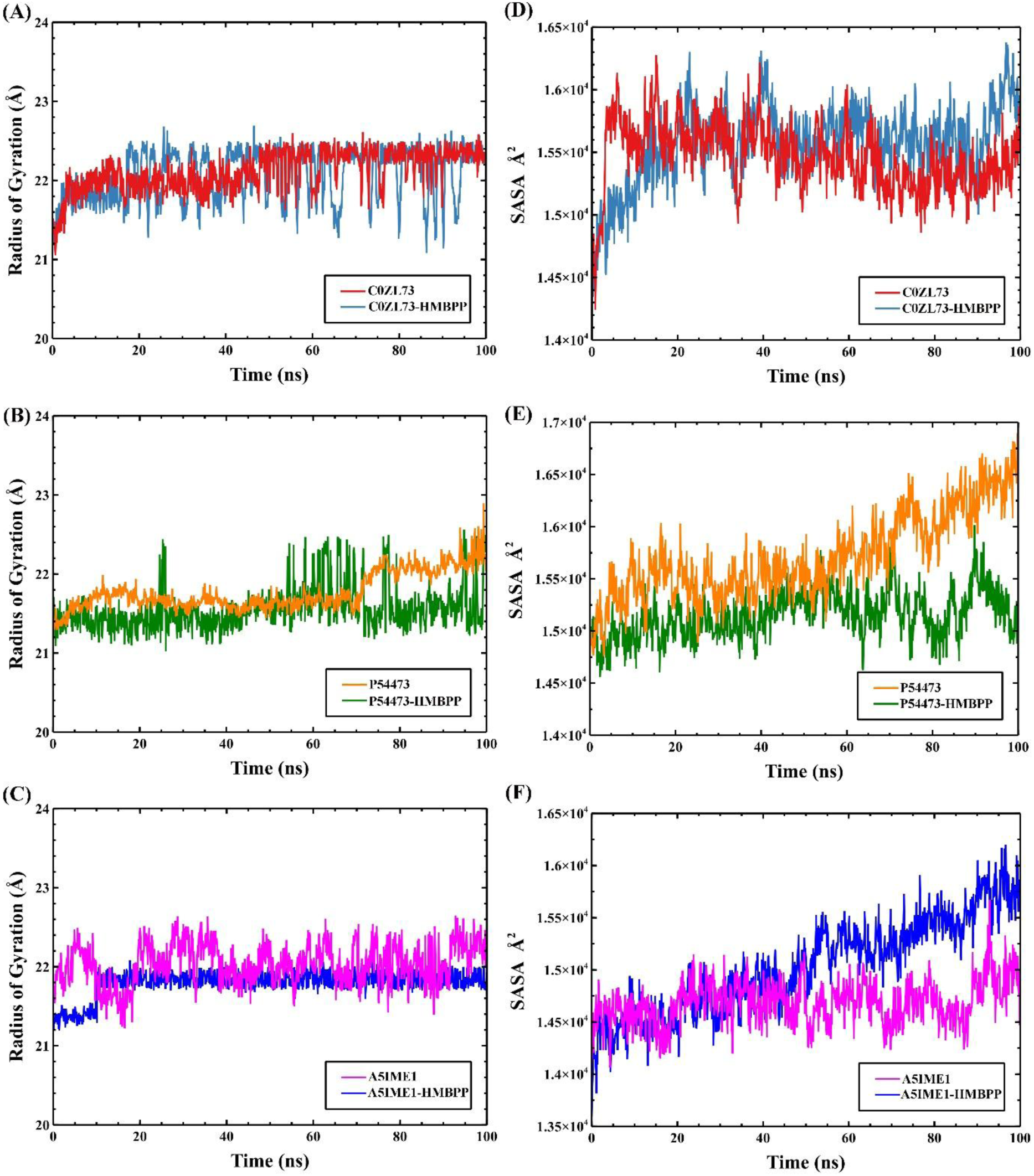
The Rg and SASA plots for the 100 ns MD simulation of the protein and protein-ligand complexes. C0ZL73-HMBPP (A, D), P54473-HMBPP (B, E), and A5IME1-HMBPP (C, F).

Comparing the Rg values of the apo form and ligand-bound form of the proteins indicates that HMBPP binding does not significantly affect the protein’s compactness and overall structure. Additionally, the stable Rg values over the simulation suggest that the proteins and their complexes with HMBPP maintain a consistent conformation^88^. The SASA values computed for the proteins and their complexes with HMBPP over 100ns are illustrated in Figures 9D-F. For C0ZL73, SASA increases slowly as the simulation progresses and then shows a decreasing trend, while for P54473 and A5IME1, SASA continues to increase slowly until the end of the simulation. The average SASA values for C0ZL73, P54473, and A5IME1 across the simulation are observed to be 15,463Å², 15,712Å², and 14,682Å², respectively. Upon attachment to the ligand, a slight decrease in average SASA value is observed in C0ZL73 (15,428 Å²) and P54473 (15,152 Å²), likely due to ligand occupancy. In the case of A5IME1, ligand attachment leads to a slight increase in average SASA value to 15,035 Å², possibly due to a slight conformational change in the protein that increases solvent exposure sites^89^.

#### 3.6.2. Intermolecular interaction between proteins and HMBPP

The intermolecular interactions between the studied proteins and HMBPP during the simulation are illustrated in Figures 10 and 11. Figure 10A shows the number of hydrogen bonds established by protein C0ZL73 and HMBPP during a 100 ns simulation. At 0ns, there are 13 hydrogen bonds, which decrease to 8 by 30 ns. Thereafter, the number of hydrogen bonds stabilizes, remaining at an average of 8 until the end of the simulation.

**Figure 10:**
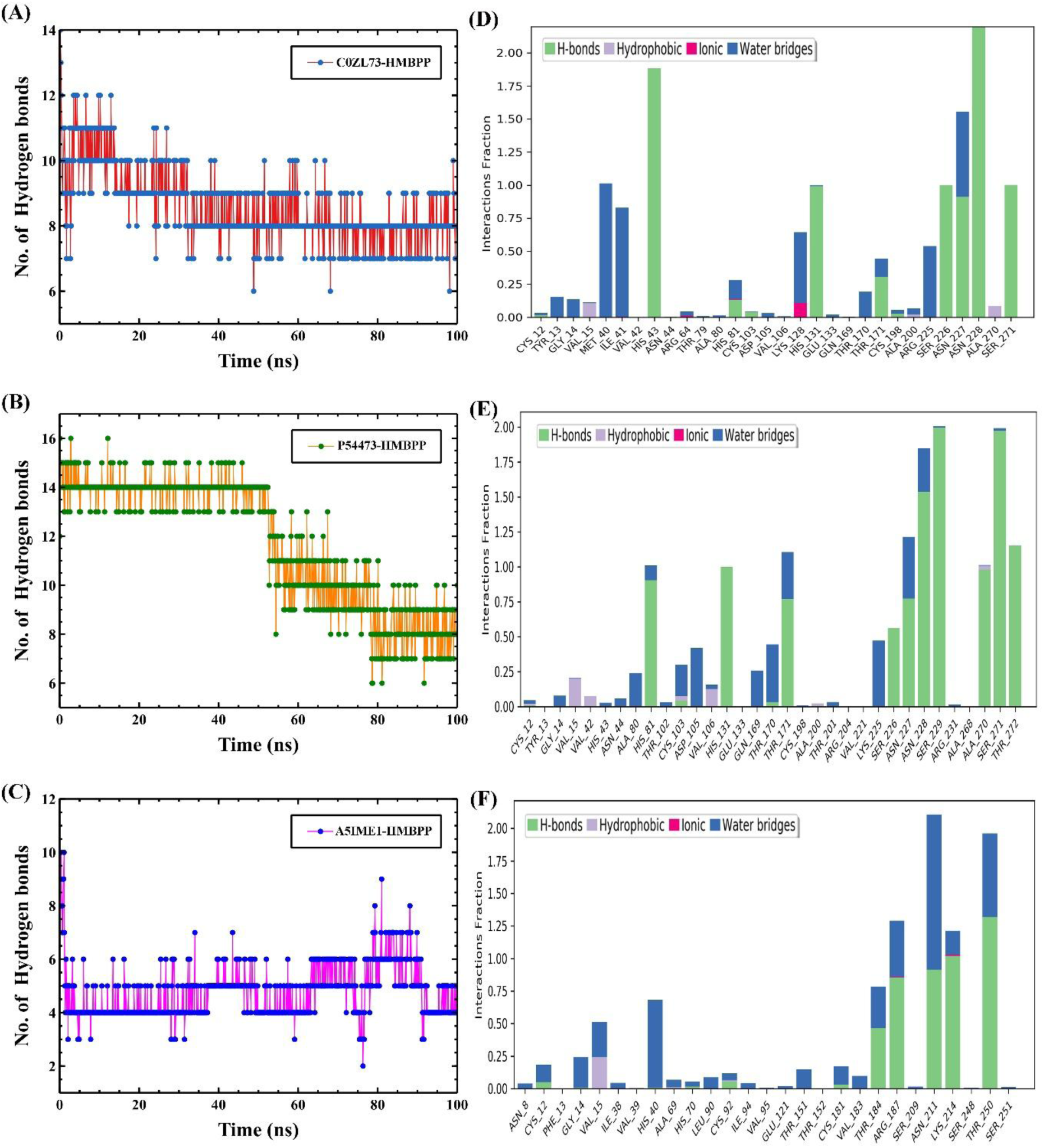
The number of hydrogen bond and fraction of interactions observed during MD simulation for C0ZL73-HMBPP (A, and D), P54473-HMBPP (B, and E), and A5IME1-HMBPP (C, and F).

**Figure 11:**
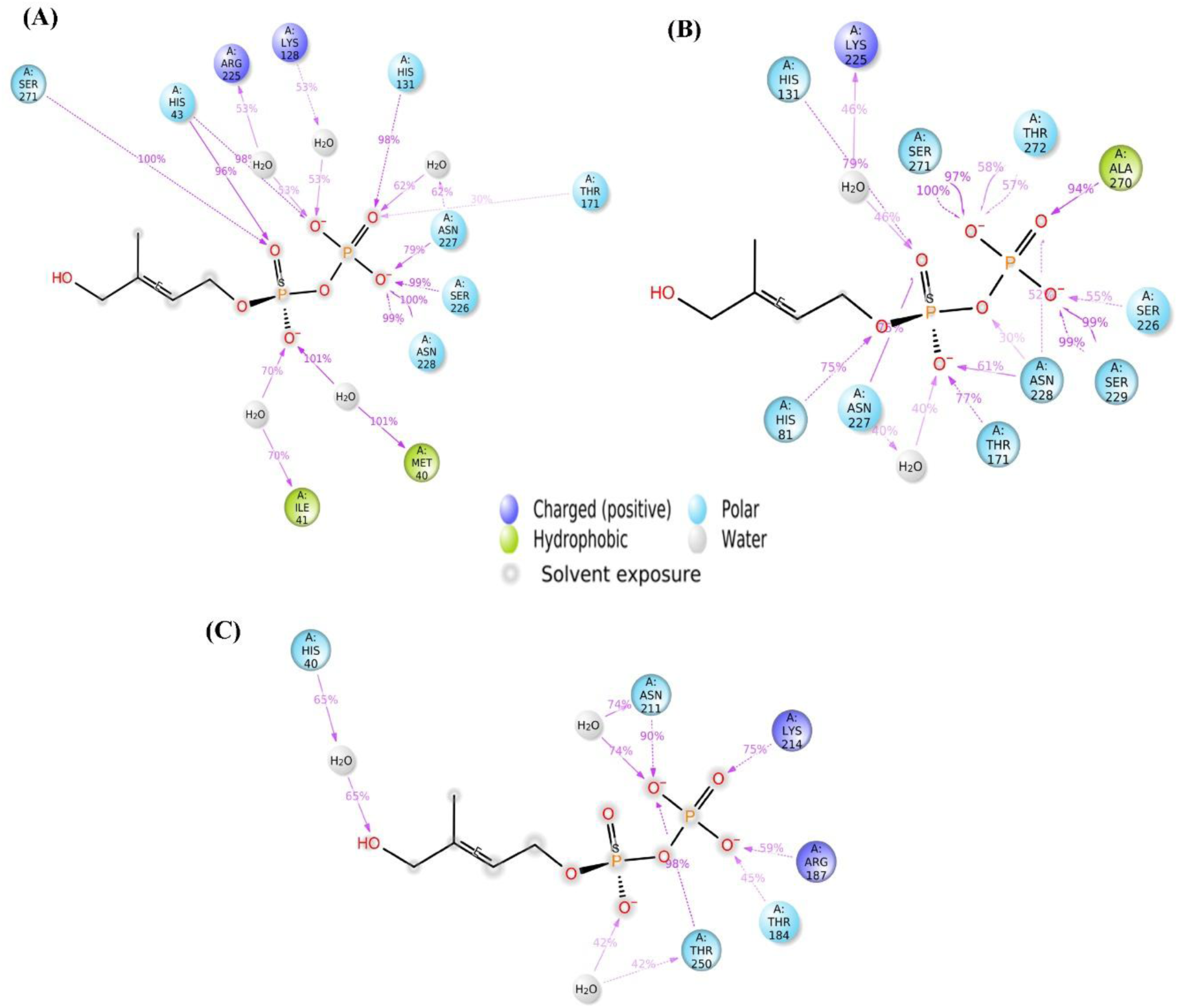
The 2D representation of fraction of various interactions observed in (A) C0ZL73-HMBPP, (B) P54473-HMBPP and (C) A5IME1-HMBPP complex.

Key residues involved in hydrogen bonding, with atleast 75% persistent interactions throughout the trajectory, or interaction fraction exceeding 0.75, include His43, His131, Ser226, Asn227, Asn228, and Ser271 (Figures 10D and 11A). For P54473-HMBPP complex (Figure 10B), the simulation starts with 12 hydrogen bonds, increasing to an average of 14 bonds until 54 ns, before gradually declining to 8 bonds at 100 ns. A set of 10 residues, viz. His81, His131, Thr171, Ser226, Asn227, Asn228, Ser229, Ala270, Ser271, and Thr272, are found involved in hydrogen bonding with HMBPP (Figures 10E and 11B). Likewise, A5IME1-HMBPP complex (Figure 10C) has 10 hydrogen bonds at 0 ns, which immediately drops to 2 and stabilizes at an average of 5 during the simulation. The residues implicated in hydrogen bonding include Arg187, Asn211, Lys214, and Thr250 (Figures 10F and 11C). Water bridges, another important interaction influencing structural stability, are present in all complexes. Water bridges involving Met40, Ile41, Lys128, Arg125, and Asn227 are found in the C0ZL73-HMBPP complex. Asp105, Gln169, Thr170, Thr171, Lys225, Asn227, and Asn228 serve as water bridges in the P54473-HMBPP complex, and likewise, the A5IME1-HMBPP complex exhibits water bridges involving His40, Thr184, Arg187, Asn211, and Thr250. Although hydrophobic and ionic interactions are also present between the proteins and the ligand, they do not persist for extended durations. Hence, C0ZL73-HMBPP and P54473-HMBPP exhibit a large number of interactions over extended periods, indicating that these complexes are more stable than the A5IME1-HMBPP complex.

## 4. Conclusion

IspH is a crucial bottleneck enzyme for enhancing the bioproduction of isopentenols. Topological analysis of the Alphafold models of the representative sequences indicate a robust structural conservation across the bacterial IspH sequences. The *B. brevis47* and *B. subtilis*168 dataset representatives C0ZL73 and P54473 exhibit a notably larger portion of conserved residues, specifically 50% and 78.98%, in contrast to their 15.28% and 7.32% of highly variable regions. The predicted interaction networks for the representative sequences show that cmk, catalyzing the bioproduction of (d)CDP from (d)CMP using ATP as a substrate, is functionally important for IspH in meeting the increased energy demand for running the DXP pathway. Despite encoding the low functionally conserved residues, only the *B. brevis*47 subgroup demonstrates the evolutionarily connected residue communities of 7- and 14-residues, demonstrating their functionally significant relevance for IspH, and *B. brevis* IspH should be experimentally investigated over *B. Subtilis*. Using the computational analysis, the study successfully extracts the key active site residues for guiding the directed evolution experiments. In addition, molecular docking and molecular dynamics simulation studies show that IspH derived from both *B. brevis*47 and *B. subtilis*168 have stable binding with HMBPP, indicating its potential to be developed into a strong mediator of the DXP pathway to enhance industrial production of isopentenols.

## Credit authorship contribution statement

Ashish Runthala: Conceptualization, Data curation, Formal analysis, Methodology, Writing – original draft; Sivudu Macherla, and Venkatramanan Varadharajan: Data curation, Investigation, Methodology. Silambarasan Tamil Selvan: Formal analysis; Manmohan Sharma and Namit Kaur: Resources, Validation; Suresh Chandra Phulara: Writing – original draft, Writing – review & editing; Noor Ahmad Shaik and Hazem K Ghneim: Writing – review & editing.

## Acknowledgements

The authors would like to thank their university for the financial support needed to complete this paper.

